# *Myh10* deficiency leads to defective extracellular matrix remodeling and pulmonary disease

**DOI:** 10.1101/414268

**Authors:** Hyun-Taek Kim, Wenguang Yin, Young-June Jin, Paolo Panza, Felix Gunawan, Beate Grohmann, Carmen Buettner, Anna M. Sokol, Jens Preussner, Stefan Guenther, Sawa Kostin, Clemens Ruppert, Aditya M. Bhagwat, Xuefei Ma, Johannes Graumann, Mario Looso, Andreas Guenther, Robert S. Adelstein, Stefan Offermanns, Didier Y.R. Stainier

## Abstract

Impaired alveolar formation and maintenance are features of many pulmonary diseases that are associated with significant morbidity and mortality. In a forward genetic screen for modulators of mouse lung development, we identified the non-muscle myosin II heavy chain gene, *Myh10*. *Myh10* mutant pups exhibit cyanosis and respiratory distress, and die shortly after birth from differentiation defects in alveolar epithelium and mesenchyme. From omics analyses and follow up studies, we find decreased Thrombospondin expression accompanied with increased matrix metalloproteinase activity in both mutant lungs and cultured mutant fibroblasts, as well as disrupted extracellular matrix (ECM) remodeling. Loss of *Myh10* specifically in mesenchymal cells results in ECM deposition defects and alveolar simplification. Notably, MYH10 expression is down-regulated in the lung of emphysema patients. Altogether, our findings reveal critical roles for *Myh10* in alveologenesis at least in part via the regulation of ECM remodeling, which may contribute to the pathogenesis of emphysema.

## Introduction

Lung development is subdivided into five chronologically and structurally distinct stages: embryonic, pseudoglandular, canalicular, saccular, and alveolarization^1,2^. During the embryonic stage, the lung bud arises from the anterior foregut endoderm. From the pseudoglandular to canalicular stages, the lung undergoes branching morphogenesis to form a tree-like tubular structure. At the saccular stage, expansion and thinning of the alveolar walls lead to a marked decrease in interstitial tissue as a prerequisite for postnatal gas exchange. Finally, the lung greatly expands its alveolar surface by alveolarization, a process by which alveolar sacs are repeatedly partitioned through septation. Alveolar sac septation is a complex process which requires interactions between lung epithelial and mesenchymal cells^3-7^. Although several studies have focused on these epithelial-mesenchymal interactions, the underlying mechanisms are still poorly understood.

Non-muscle myosin II (NM II) plays fundamental roles in the maintenance of cell morphology, cell adhesion and migration, as well as cell division^8-10^. NM II molecules consist of three peptide pairs: a pair of NM II heavy chains (NMHC II), a pair of regulatory light chains (RLCs), and a pair of essential light chains (ELCs). In mammals, three different genes, *Myosin heavy chain 9* (*MYH9*), *MYH10*, and *MYH14,* encode the NMHC II proteins, NMHC IIA, NMHC IIB, and NMHC IIC, respectively^11,12^. NM II proteins are highly expressed during lung development^13^, and their inhibition using blebbistatin leads to defects in branching morphogenesis and epithelial cell shape and orientation^14^. To date, however, genetic studies of NM II genes during lung development and homeostasis have not been conducted.

Many pulmonary diseases including chronic obstructive pulmonary disease (COPD)^15,16^ and interstitial lung disease (ILD)^17^ are associated with significant morbidity and mortality due to impaired alveolar formation and maintenance. Emphysema is one form of COPD that results from the enzymatic destruction of lung ECM components, including elastin and collagen, thereby leading to destruction of alveolar walls and airspace enlargement^15,16,18^. Recently, several animal models have been established to study the pathophysiology of emphysema, including exposure to cigarette smoke and protease instillation, as well as genetic manipulation^18-20^. The pathophysiological role of NM II in pulmonary emphysema, however, has not been investigated.

Here, starting with a forward genetic screen in mouse, we reveal an unexpected role for the NM II-associated actomyosin network in ECM remodeling during lung development and disease. Notably, our results from animal models and human patients suggest that alterations of the actomyosin network by loss of *MYH10* function contribute to the pathogenesis of emphysema and may provide a promising target for preventive care of emphysema.

## Results

### Identification of lung mutants following ENU mutagenesis

To identify novel factors regulating early postnatal lung development in mouse, we carried out a forward genetic screen using ENU mutagenesis^21^. From screening the F3 progeny of 170 G1 mice, we identified 8 families exhibiting a recessive alveolar collapse phenotype. Mutant newborns from one of these families were cyanotic and exhibited respiratory distress with full penetrance (Fig. 1a, Supplementary Movie 1). As their lungs did not inflate, mutant pups died within 24 hours after birth (Fig. 1b, c). Histological and morphometric analyses revealed reduced alveolar spaces and thick alveolar septal walls (Fig. 1d), along with dilated cardiomyopathy and hydrocephalus^22^ (Supplementary Fig. 1a, b). We observed no histological differences between wild-type and mutant littermate lungs at the pseudoglandular stage. However, from embryonic day (E) 17.5 onward, the mutant lungs displayed reduced saccular spaces and more abundant mesenchyme compared to wild-type siblings (Fig. 1e).

**Figure 1.**
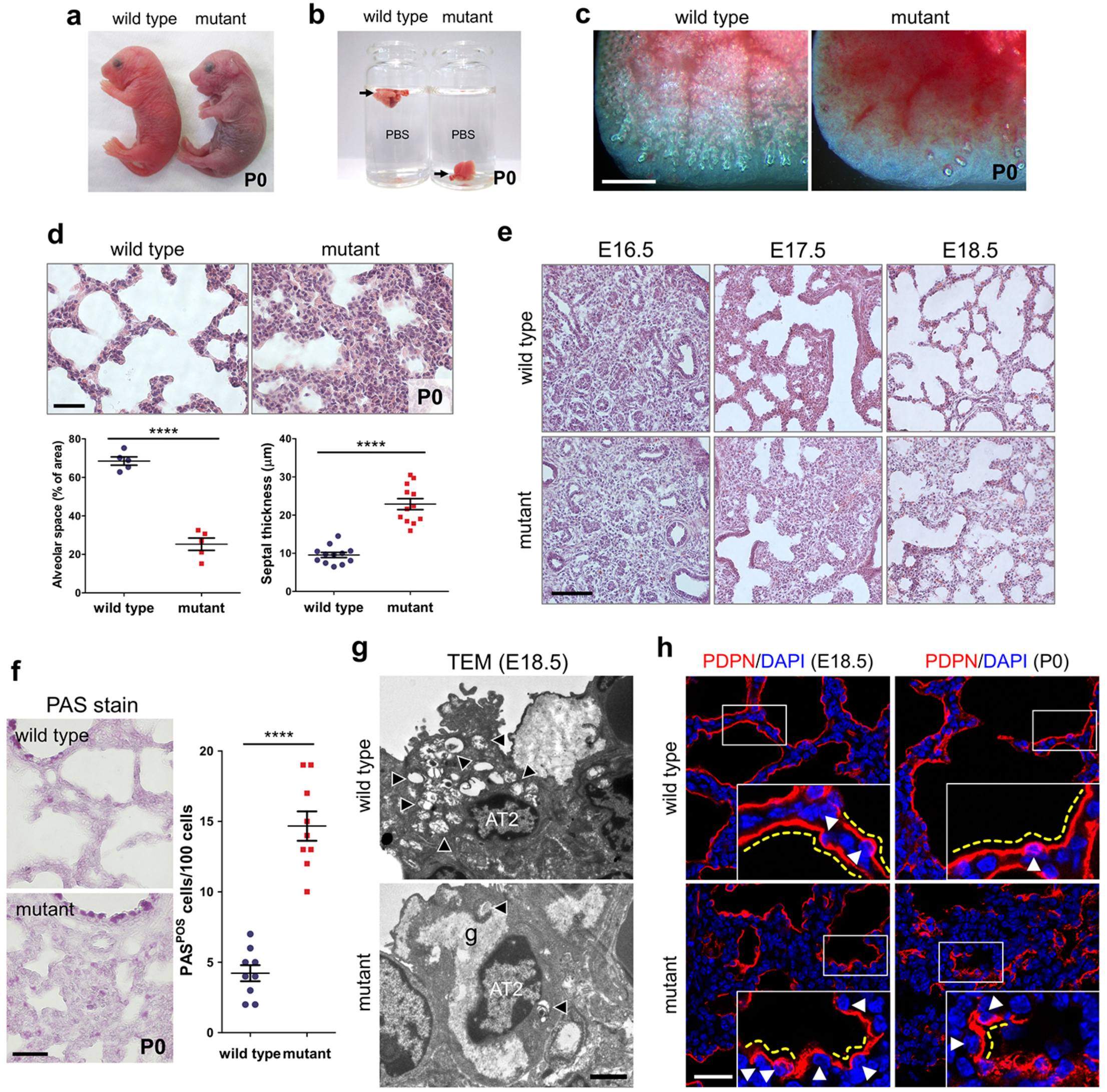
The ENU-induced mutant exhibits lung development defects. **(a)** Gross morphology of wild-type (n=15) and mutant (n=12) P0 mice. (**b)** Floating assay for wild-type (n=10) and mutant (n=8) P0 lungs. **(c)** Representative pictures of distal lung from wild-type (n=10) and mutant (n=8) P0 mice. **(d)** Hematoxylin and Eosin (H&E) staining and morphometric analysis of alveolar space and septal thickness in wild-type (n=15) and mutant (n=10) P0 lungs. **(e)** H&E staining of wild-type (n=6 for each stage) and mutant (n=5 for each stage) lungs at E16.5, E17.5, and E18.5. **(f)** PAS staining (n=15 wild types, n=10 mutants) and phenotype quantification at P0. **(g)** Representative TEM images of AT2 cells in wild-type (n=3) and mutant (n=3) lungs at E18.5. Arrowheads point to lamellar bodies in AT2 cells; g, glycogen. **(h)** Immunostaining for PDPN in wild-type (n=8 per each stage) and mutant (n=8 per each stage) lungs at E18.5 and P0. Arrowheads point to AT1 nuclei; yellow dotted lines outline AT1 cell morphology. Error bars are means ±s.e.m. \*\*\*\**P* < 0.0001, two-tailed Student’s *t* test. Scale bars: 500 μm (c), 50 μm (d, e), 30 μm (f, h), 2 μm (g).

To investigate whether the increased tissue density observed in the mutant lungs resulted from increased cell proliferation, reduced apoptosis, or differentiation defects, we performed immunostaining using two proliferation markers, PCNA and phospho-histone 3 (pH3), and quantified positive cells from E17 to postnatal day 0 (P0). To differentiate between mesenchymal and epithelial cells, we co-stained for the epithelial cell markers NKX2-1 and E-cadherin (E-cad). Interestingly, both epithelial and mesenchymal cells exhibited increased cell proliferation in mutant lungs compared to wild-type siblings (Supplementary Fig. 1c, d, 2a, b), explaining at least in part the observed increase in tissue density. A minor increase in the number of apoptotic cells was also observed in mutant lungs compared to wild-type siblings (Supplementary Fig. 1c, 2c), suggesting that loss of mutant cells does not contribute in a major way to the observed phenotypes.

Next, we examined the differentiation status of mutant epithelial and mesenchymal cells by staining for several marker proteins. Interestingly, while the number of Vimentin-positive fibroblasts^23^ was dramatically decreased in mutant lungs, unaltered NG2-positive pericyte-like cells^23^ and endothelial cell numbers were observed in mutant lungs compared to wild-type siblings (Supplementary Fig. 1d). Moreover, mutant lungs exhibited an increase in the number of AT2 (alveolar epithelial type II) cells and AT2 cell marker (*Sftpa, Sftpc, Abca3*) expression levels (Supplementary Fig. 1e, 2d, e). Concurrently, we found a marked decrease in the expression levels of AT1 (alveolar epithelial type I) cell markers (HOPX, Podoplanin (PDPN), and RAGE) ^24^ as well as of a mature AT2 cell marker (LAMP3)^25^ (Supplementary Fig. 1e, f, 2e).

Immature AT2 cells store glycogen which is converted into surfactant in lamellar bodies as these cells differentiate^26^. To examine the differentiation state of AT2 cells, we performed Periodic acid-Schiff (PAS) staining, which revealed a substantial increase in the number of PAS-positive AT2 cells in P0 mutant lungs (Fig. 1f). By transmission electron microscopy (TEM), wild-type AT2 cells contained numerous lamellar bodies and apical microvilli at E18.5, whereas mutant AT2 cells displayed abundant glycogen, but fewer lamellar bodies and microvilli, indicating that they failed to differentiate (Fig. 1g). In addition, whereas AT1 cells in wild-type lungs covered the alveolar surface and exhibited a flattened morphology at E18.5 and P0, most AT1 cells in mutant lungs appeared densely packed and failed to flatten out, as observed by PDPN immunostaining (Fig. 1h). The numbers of distal airway epithelial cells including club cells and ciliated cells did not appear to be significantly altered between mutants and wild-type siblings (Supplementary Fig. 1g). Taken together, these results indicate that mutants exhibit disrupted lung epithelial and mesenchymal cell differentiation, resulting in abnormal AT1 cell morphology and defective saccular development.

### Identification of a missense mutation in the *Myh10* gene

To identify the causative mutation, we carried out whole-exome sequencing of mutant and wild-type siblings and identified a missense Leu-to-Arg mutation (c.T1373G;p.L458R) in the motor domain of NMHC II-B, which is encoded by the *Myh10* gene (Fig. 2a, d). Next, we performed genetic linkage analysis by genotyping 102 G4 and G5 mutant mice and found complete association between the lung phenotype and the *Myh10* mutation (Fig. 2b, c). We then performed a complementation test between the ENU-induced *Myh10* mutant and a *Myh10* null allele^22^. Trans-heterozygous mice exhibited the same lung and heart phenotypes as the *Myh10* global^27,28^ and ENU-induced mutants, indicating that the loss of *Myh10* function is responsible for the lung phenotype (Supplementary Fig. 3a, b). In addition, the localization of MYH10^L458R^ was different than that of MYH10^WT^ in cultured fibroblasts: while MYH10^WT^-Myc was localized as scattered punctae in stress fibers and lamellae^29^, MYH10^L458R^-Myc appeared dissociated from the actin bundles of stress fibers (Supplementary Fig. 3c). To quantify the expression levels of *Myh10* mRNA and protein in the P0 mutant lungs, we used RT-qPCR, western blotting, and immunostaining. Expression of *Myh10* mRNA was not significantly altered in *Myh10* ^*L458R/L458R*^ (hereafter *Myh10*^*-/-*^) lungs compared to *Myh10*^*+/+*^ and *Myh10*^*+/L458R*^ (hereafter *Myh10*^*+/-*^) lungs. However, MYH10 protein expression was dramatically reduced in *Myh10* lungs compared to *Myh10* and *Myh10* lungs (Supplementary Fig. 3d). These data suggest that possibly due to their inability to bind actin bundles, MYH10^L458R^ proteins are highly unstable and thus subsequently degraded. To test the expression pattern of *Myh10* in the developing lung, we performed *in situ* hybridization and immunostaining. At E13.5, *Myh10* transcripts were specifically detected in mesenchymal tissue in both whole-mount and cryosectioned lungs (Fig. 2e). At E16.5, *Tbx4*-positive mesenchymal cells^30^ co-expressed MYH10, whereas E-cadherin-positive epithelial cells did not (Fig. 2f, g). At E18.5, MYH10 expression co-localized with α-SMA and PDGFR-β expression in the lung mesenchyme, but not with NG2, VE-cadherin, or PDPN expression (Fig. 2g, Supplementary Fig. 3e). These results indicate that in developing lungs, *Myh10* is specifically expressed in mesenchymal cells including myofibroblasts, lipofibroblasts, and smooth muscle cells but not in endothelial cells, pericyte-like cells, or epithelial cells.

**Figure 2.**
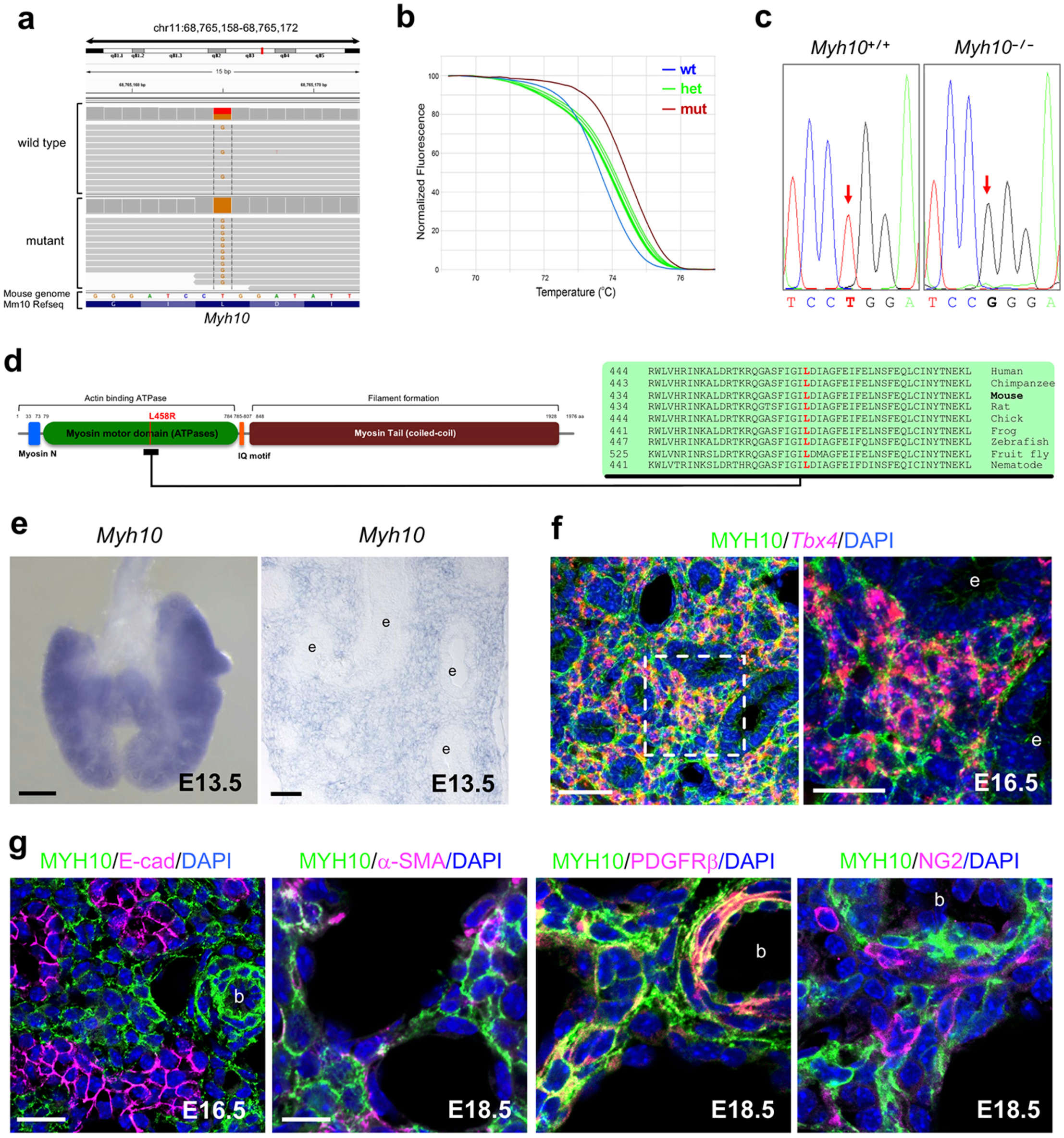
Isolation of the causative lesion and expression pattern of *Myh10*. (**a)** Integrative genomics viewer (IGV) snapshot of the c.T1373G mutation in *Myh10* (chr 11:68,765,165). **(b)** Normalized melting curves showing the *Myh10* mutation by high resolution melting analysis (HRMA). **(c)** Chromatogram of two different genotypes of *Myh10* by Sanger sequencing. **(d)** Schematic diagram of MYH10 protein domains and the relative position of the L458 residue which is conserved from worms to humans. **(e)** *In situ* hybridization for *Myh10* expression in E13.5 whole-mount and on cryosectioned lungs; e, epithelium. **(f)** Double staining for *Tbx4* mRNA (red) and MYH10 protein (green) on E16.5 lung sections; e, epithelium. **(g)** Immunostaining for MYH10, E-cadherin (marking epithelial cells), α-SMA (marking myofibroblasts and smooth muscle cells), PDGFR-β (marking myofibroblasts and smooth muscle cells), and NG2 (marking pericyte-like cells) on E16.5 and E18.5 lung sections; b, blood vessel. Scale bars: 200 μm (e (left)), 50 μm (e (right), f (left)), 20 μm (f (right), g (left)), 10 μm (g (right)).

### *Myh10* deficiency causes defects in ECM remodeling

To understand the molecular mechanisms of *Myh10* function during lung development, we analyzed the transcriptome of *Myh10*^*+/+*^ and *Myh10*^*-/-*^ lungs at the saccular stage. Many of the genes downregulated in *Myh10*^*-/-*^ lungs are known to participate in cell adhesion and contraction, as well as ECM remodeling (Supplementary Fig. 4a, Dataset 1), prompting us to investigate the roles of NM II/MYH10 in this process. First, we examined the expression levels of several ECM components. Immunostaining and western blot data showed that expression of Fibronectin (FN), type I Collagen (COL I), and Tropoelastin, which form the interstitial ECM^31^, appeared significantly reduced in E18.5 *Myh10*^*-/-*^ lungs (Fig. 3a, Supplementary Fig. 4b). In contrast, expression of basement membrane (BM) ECM components^31^ type II (COL II) and type IV Collagen (COL IV), as well as that of Laminin, were not altered, suggesting that defects in MYH10 function affect the expression of proteins found in the interstitial ECM of alveoli but not of those in the alveolar BM (Supplementary Fig. 4d). However, RNA expression of *FN, Col I*, and *Tropoelastin* was unaltered in E18.5 *Myh10*^*-/-*^ lungs compared to wild-type lungs (Supplementary Fig. 4c). Using TEM, we analyzed ECM organization in the alveolar interstitium at E18.5. Mature elastic fibers were largely absent in the ECM of *Myh10*^*-/-*^lungs, and elastic bundles appeared sparse and fragmented compared to the wild-type pattern (Fig. 3b). These data suggest that *Myh10* deficiency disrupts expression levels and localization of ECM components, including FN, Col I, and Elastin. MYH10 is known to mediate primary cilia formation in a human retinal pigmented epithelial (RPE1) cell line^32, 33^. To test whether MYH10 regulates cilia formation in embryonic fibroblasts, we performed immunostaining for anti-acetylated α-tubulin to detect primary cilia. We found no obvious defects in cilia formation in *Myh10* deficient fibroblasts compared to wild-type fibroblasts (Supplementary Fig. 4e). Moreover, expression of components of the Wnt, Sonic hedgehog/Shh, and Fibroblast growth factor/FGF signaling pathways, which are involved in lung cell proliferation and differentiation^4, 34^, was not altered in *Myh10*^*-/-*^lungs compared to wild-type siblings (Supplementary Fig. 4f), indicating that these signaling pathways are not responsible for the phenotypes observed in *Myh10* deficient fibroblasts and lungs.

**Figure 3.**
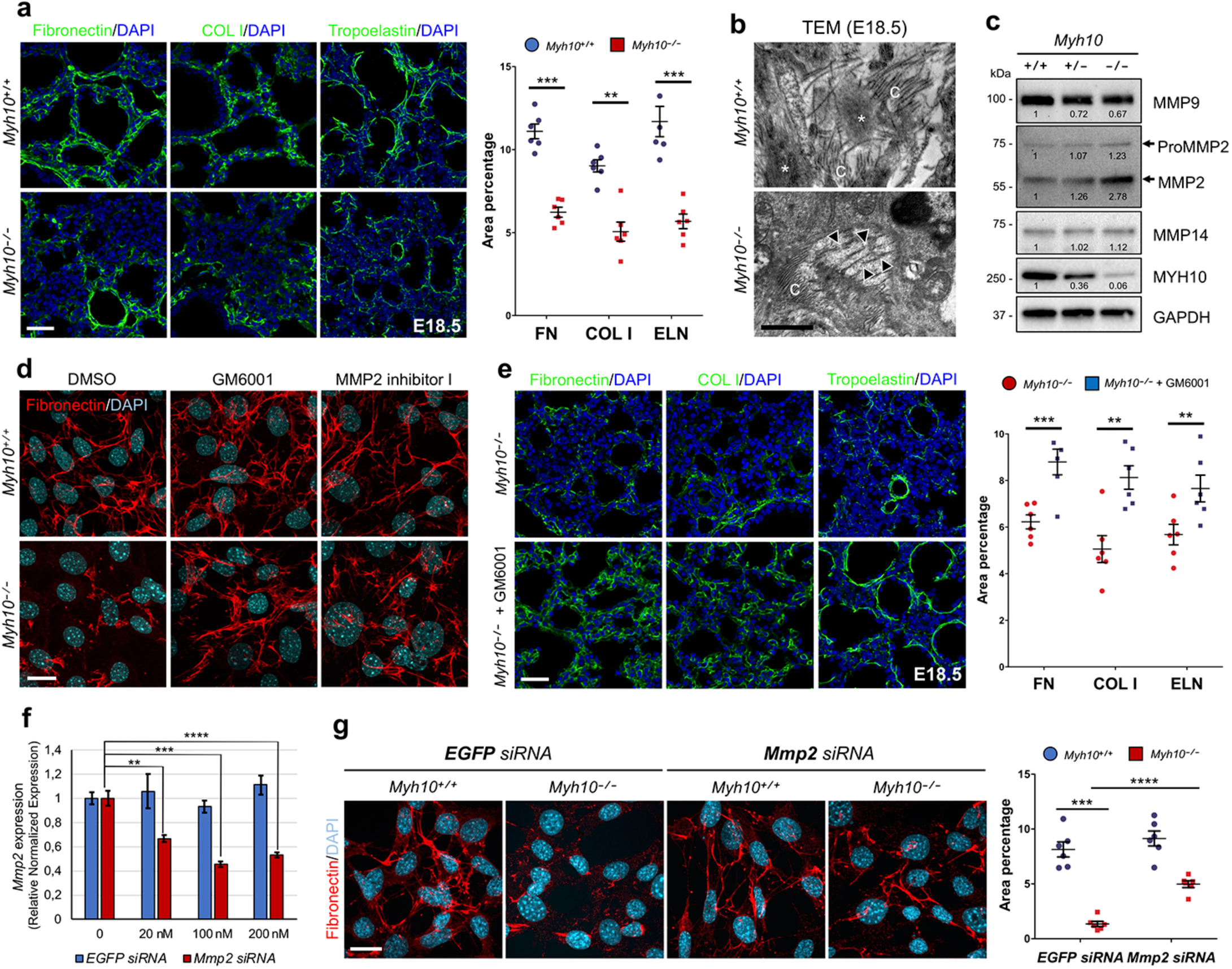
*Myh10* deficiency leads to disrupted ECM remodeling. **(a)** Immunostaining and quantification for Fibronectin, Type I collagen (COL I), and Tropoelastin on *Myh10*^+/+^ (n= 10) and *Myh10*^*-/-*^ (n=10) E18.5 lung sections. **(b)** Representative TEM images comparing elastic fibers in *Myh10*^+/+^ (n=3) and *Myh10*^*-/-*^ (n=3) E18.5 lungs. Asterisks and arrowheads point to bundles and fragmented bundles of elastic fibers, respectively; c, collagenous fibrils. **(c)** Representative western blot (from three individual sets of lung lysates) of *Myh10*^+/+^, *Myh10*^+/-^, and *Myh10*^*-/-*^ lung lysates at E18.5. Values represent the densitometric ratio of *Myh10*^+/-^ and *Myh10*^*-/-*^ to *Myh10*^+/+^ after normalization to GAPDH. **(d)** Immunostaining (from five individual sets of cultured cells) for Fibronectin in DMSO, GM6001, or MMP2 inhibitor I-treated *Myh10*^+/+^ and *Myh10*^-/-^ embryonic fibroblasts. **(e)** Immunostaining and quantification for Fibronectin, COL I, and Tropoelastin on *Myh10*^*-/-*^ (n=5) and GM6001-injected *Myh10*^*-/-*^ (n=5) E18.5 lung sections. **(f)** Expression level of *Mmp2* in *EGFP* siRNA and *Mmp2* siRNA-treated *Myh10* embryonic fibroblasts by RT-qPCR. **(g)** Immunostaining and quantification for Fibronectin in *EGFP* siRNA and *Mmp2* siRNA-treated *Myh10* and *Myh10* embryonic fibroblasts. Error bars are means ±s.e.m. \*\**P* < 0.01, \*\*\**P* < 0.001, \*\*\*\**P* < 0.0001, two-tailed Student’s *t* test. Scale bars: 30 μm (a, e), 20 μm (d, g), 500 nm (b).

### Excess MMP2 activity causes defective ECM remodeling

Metalloproteinases are a group of proteins essential for ECM remodeling, which includes members of the matrix metalloproteinase (MMP) and a disintegrin and metalloproteinase with thrombospondin motifs (ADAMTS) family members^35, 36, 37^. During embryonic development, MMP-2 is one of the most highly expressed metalloproteinases in the lung^38^. Through both MMP activity assays and western blotting, we found a dramatic increase of MMP-2 activity in lysates of E18.5 *Myh10*^*-/-*^ lungs and of *Myh10*^*-/-*^ cultured fibroblasts compared to wild-type samples (Fig. 3c, Supplementary Fig. 5a, b). To test whether MMP inhibition could rescue the *Myh10* mutant phenotypes, we first treated wild-type and *Myh10* mutant embryonic fibroblasts^39^ with GM6001, a broad-spectrum MMP inhibitor^40^, and with the MMP2-specific inhibitor I^41^. Wild-type fibroblasts displayed well-organized, linear FN fibers; in contrast, only small aggregates of FN marked the cell surface of *Myh10* deficient fibroblasts. Interestingly, the FN matrix defect was partially rescued by treatment with GM6001 or MMP2 inhibitor I (Fig. 3d, Supplementary Fig. 5c). Next, we administered GM6001 to pregnant females daily, covering a developmental time window from E16.5 to E18.5. This regimen was sufficient to partially rescue defective ECM remodeling in *Myh10*^*-/-*^ lungs (Fig. 3e). In addition, the FN matrix defect in *Myh10*^*-/-*^ fibroblasts was partially rescued by siRNA-mediated *Mmp2* knock-down (Fig. 3f, g). To investigate at which specific step *Myh10* regulates ECM production, we assessed the expression of elastin assembly components in *Myh10* deficient lungs. The expression of *Fibrillin-1*/*Fbn1, Fibrillin-2*/*Fbn2, Fibulin-4*/*Fbln4, Fibulin-5*/*Fbln5, Lox*, and *Loxl1*^42,43^ was not significantly changed in E18.5 *Myh10* lungs compared to wild-type lungs (Supplementary Fig. 5d). Moreover, *Myh10* deficiency did not cause defects in FN assembly in fibroblasts (Supplementary Fig. 5e), altogether suggesting that *Myh10* does not directly control the expression, crosslinking, or assembly of ECM components in fibroblasts or developing lungs. Collectively, these results suggest that *Myh10* deficiency leads to increased MMP activity, which disrupts ECM remodeling in fibroblasts *in vitro* and in developing lungs *in vivo.*

To understand how MYH10 regulates MMP activation, we performed proteomic profiling of wild-type and mutant fibroblasts and detected lower levels of Thrombospondins (THBS1 and THBS2) in *Myh10*^*-/-*^fibroblasts (Fig. 4a and b, Supplementary Dataset 2). THBSs are matricellular proteins that interact with MMPs to inhibit their activities^44-46^. Consistently, we found a dramatic decrease in THBS1 and THBS2 expression levels in cell lysates and conditioned media of mutant fibroblasts as well as in *Myh10*^*-/-*^lungs (Fig. 4c, Supplementary Fig. 5f, g), further validating our proteomics results. Importantly, overexpression of *Thbs2* partially rescued the FN matrix defect in *Myh10*^*-/-*^fibroblasts (Fig. 4d), indicating that MYH10 regulates MMP activation through THBS.

**Figure 4.**
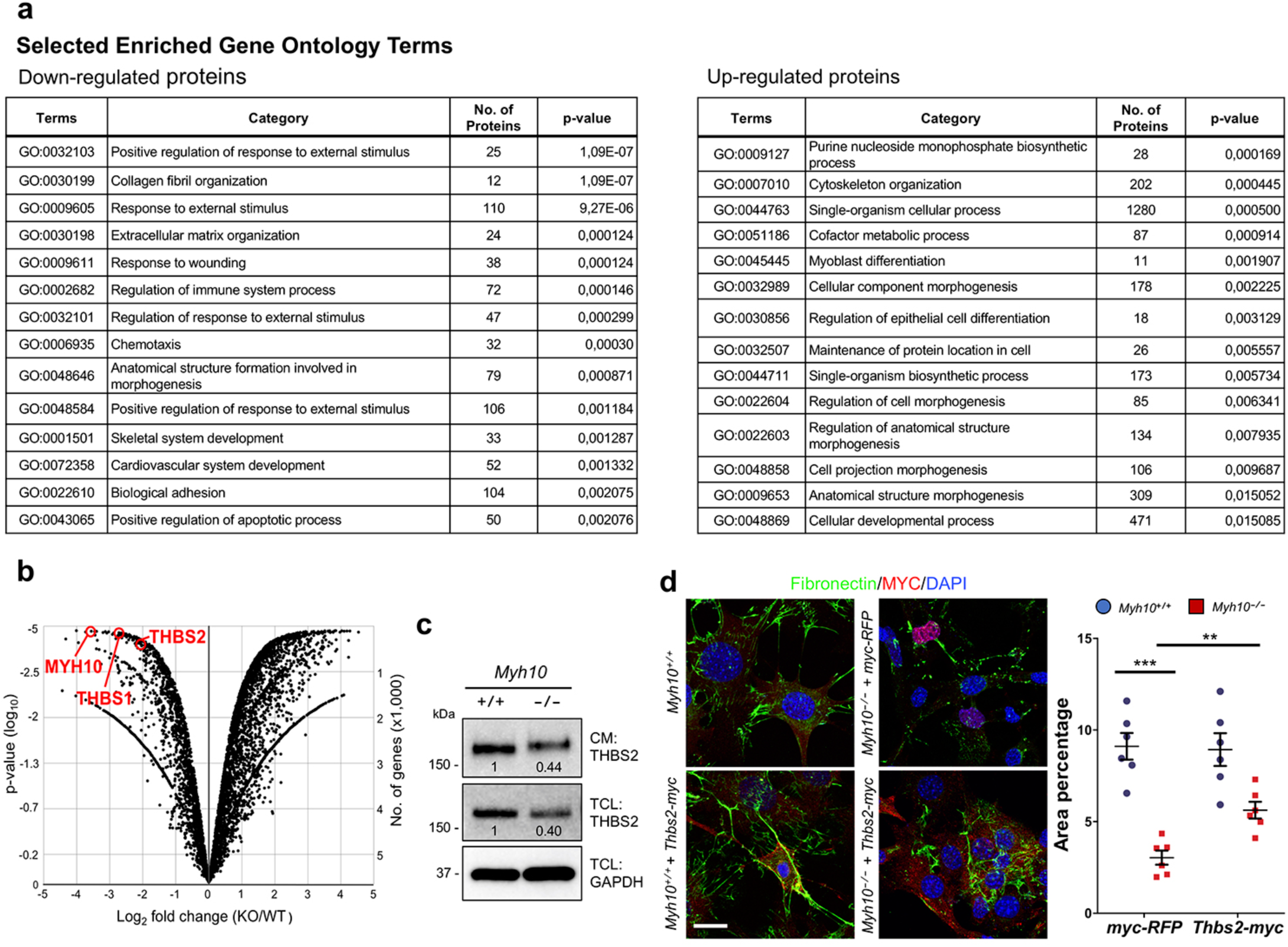
MYH10 modulates MMP activity by regulating THBS. **(a)** GO classification (all components with *p-value* <0.05) of differentially expressed proteins between *Myh10*^+/+^and *Myh10*^-/-^fibroblasts. **(b)** Volcano plot of the log_2_ ratio of differentially expressed proteins between *Myh10*^+/+^(n=3) and *Myh10*^-/-^(n=3) fibroblasts. **(c)** Representative western blot (from three individual sets of cultured cells) for THBS2 from *Myh10*^+/+^ and *Myh10*^-/-^fibroblasts. Values represent the densitometric ratio of *Myh10* ^-/-^to *Myh10*^+/+^ after normalization to GAPDH. CM, conditioned medium; TCL, total cell lysate. **(d)** Immunostaining and quantification (from four individual sets of cultured cells) for Fibronectin and Myc epitope of *Myh10*^+/+^and *Myh10*^-/-^ fibroblasts transfected with *myc-RFP* or *Thbs2-myc*. Error bars are means ±s.e.m. \*\**P* < 0.01, \*\*\**P* < 0.001, two-tailed Student’s *t* test. Scale bar: 20 μm (d).

### *Myh10* function is required in lung mesenchymal cells

To determine which cells require *Myh10* function during mouse lung development, we utilized a floxed *Myh10*^*fl/fl*^line^47^. First, we deleted *Myh10* expression in lung epithelial or endothelial cells by using *Shh-Cre* or *Tek-CreER*^*T2*^, respectively, and found no obvious breathing dysfunction or lung defects (Supplementary Fig. 6a-g). To test *Myh10* function in the lung mesenchyme, we crossed *Gli1-CreER*^*T2*^with *Myh10*^*flox*^mice (hereafter *Gli1*-*Myh10*^*cKO*^). By injecting tamoxifen into *Gli1*-*Myh10*^*cKO*^;*tdTomato* mice at E13.5-E15.5, we found that *Gli1*-positive cells did not overlap with E-cadherin-expressing epithelial cells in P0 lungs (Fig. 5a). *Gli1*-*Myh10*^*cKO*^ newborn pups exhibited respiratory distress and cyanosis (Fig. 5b). Floating lung assays from wild-type and *Gli1*-*Myh10*^*cKO*^ newborns showed that the mutant lungs failed to inflate with air at birth (Fig. 5c, d). The expression of *Myh10* mRNA and protein was dramatically decreased in *Gli1*-*Myh10*^*cKO*^ lungs as compared to wild-type and heterozygous littermates (Supplementary Fig. 7a, b). *Gli1*-*Myh10*^*cKO*^ mice also exhibited alveolar collapse, with no apparent cardiac defects (Fig. 5e, Supplementary Fig. 7c). *Gli1*-*Myh10*^*cKO*^ lungs exhibited an increased number of proliferating epithelial (NKX2-1/PCNA double positive) and mesenchymal (NKX2-1-negative/PCNA-positive) cells, as well as an increased number of AT2 cells (SFTPC-positive), as clearly in the ENU-induced allele (Fig. 5f, g). Finally, *Gli1*-*Myh10*^*cKO*^ lungs exhibited clearly decreased FN staining intensity, as well as truncated and disorganized elastin fibers (Fig. 5h). In summary, mesenchyme-specific loss of *Myh10* led to lung defects that were observed in the global *Myh10* knockout as well as in the ENU-induced *Myh10*^*L458R*^allele. These results indicate that *Myh10* is specifically required in lung mesenchyme to promote ECM remodeling and cell differentiation.

**Figure 5.**
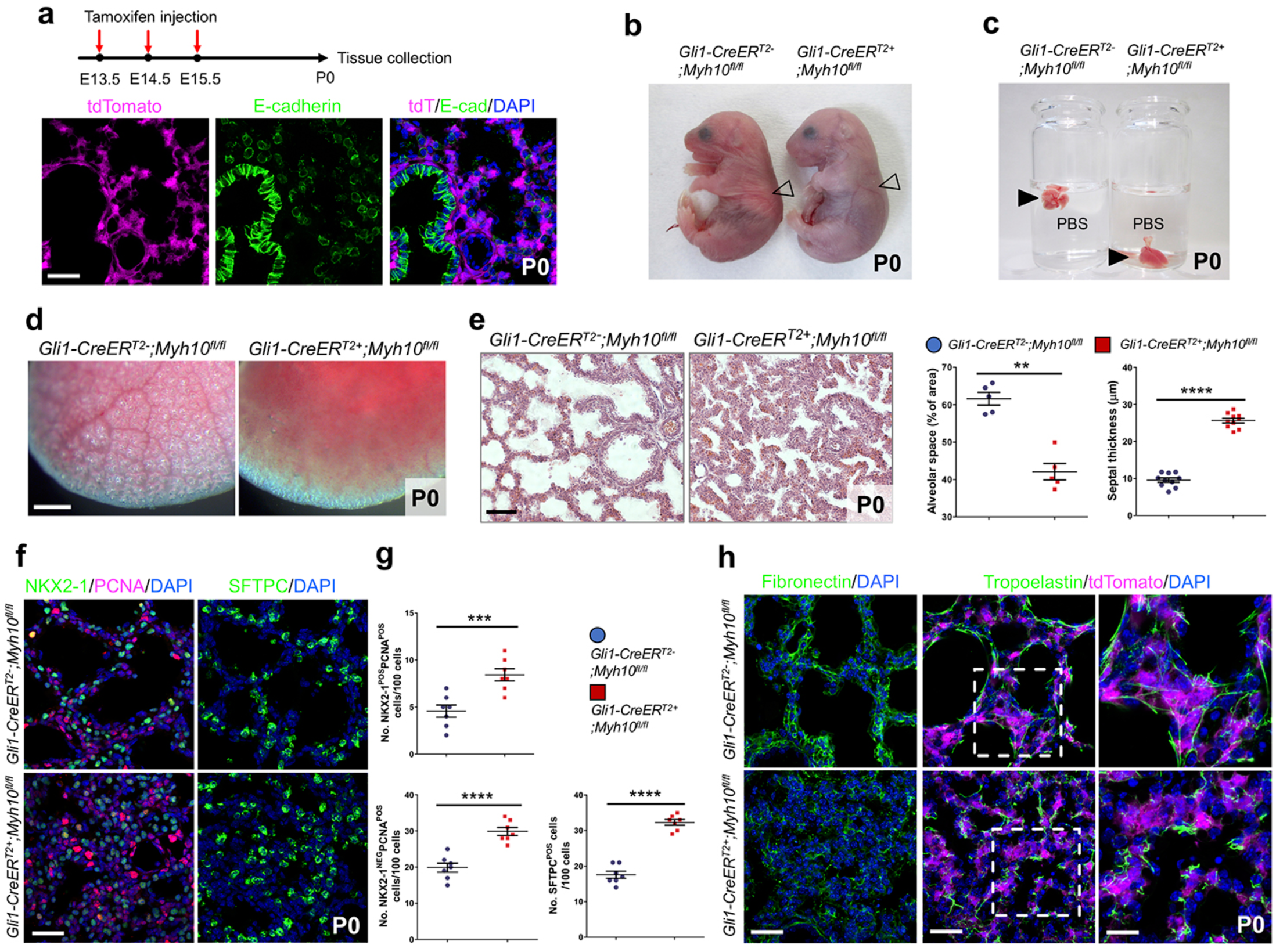
Mesenchyme specific *Myh10* deletion disrupts lung development. **(a)** Diagram indicating time points of tamoxifen injections and tissue collection. Immunostaining for E-cadherin in *Gli1-Cre*^*ERT2*^;*ROSA26*^*tdTomato*^P0 lungs. **(b)** Gross morphology of wild-type (n=10) and *Gli1-Myh10*^*cKO*^ (n=8) P0 mice. Arrowheads point to inflated (left) and non-inflated lungs (right). (**c)** Floating assay for wild-type (n=10) and *Gli1-Myh10*^*cKO*^ (n=8) P0 lungs. **(d)** Representative pictures of distal lung from wild-type (n=10) and *Gli1-Myh10*^*cKO*^ (n=8) P0 mice. **(e)** H&E staining and morphometric analysis of alveolar space and septal thickness in wild-type (n=10) and *Gli1-Myh10*^*cKO*^ (n=8) lungs at P0. **(f)** Immunostaining for NKX2-1, PCNA, and SFTPC in wild-type (n=10) and *Gli1-Myh10*^*cKO*^ (n=8) P0 lungs. **(g)** Quantification of cell proliferation and differentiation in wild-type and *Gli1-Myh10*^*cKO*^ P0 lungs. **(h)** Immunostaining for Fibronectin, Tropoelastin, and tdTomato in wild-type (n=10) and *Gli1-Myh10*^*cKO*^ (n=8) P0 lungs. Error bars are means ±s.e.m. \*\**P* < 0.01, \*\*\**P* < 0.001, \*\*\*\**P* < 0.0001, two-tailed Student’s *t* test. Scale bars: 500 μm (d), 50μm (e), 30 μm (a, f, h (left, middle)), 15 μm (h (right)).

### *Myh10* is required for lung alveologenesis

As MYH10 is still highly expressed in mesenchymal tissues throughout postnatal and adult stages (Supplementary Fig. 3e), we investigated *Myh10* function during lung maturation and homeostasis. To test the role of *Myh10* in alveologenesis, we deleted *Myh10* in mesenchymal cells at early postnatal stages before secondary septae formation^1,2,5^ (Supplementary Fig. 7d). By injecting tamoxifen into *Gli1*-*Myh10*^*cKO*^;*tdTomato* mice at P0, we found that tdTomato-expressing cells did not overlap with E-cadherin-expressing epithelial cells in P7 lungs (Supplementary Fig. 7d-f). *Gli1*-*Myh10*^*cKO*^ mice displayed a simplified, dilated alveolar structure and shorter secondary septae compared to wild-type siblings (Fig. 6a, b). Furthermore, FN expression was significantly decreased, and elastin fibers appeared disorganized and truncated (Fig. 6c), suggesting that *Myh10* is also required for secondary septation through ECM remodeling during postnatal lung development.

**Figure 6.**
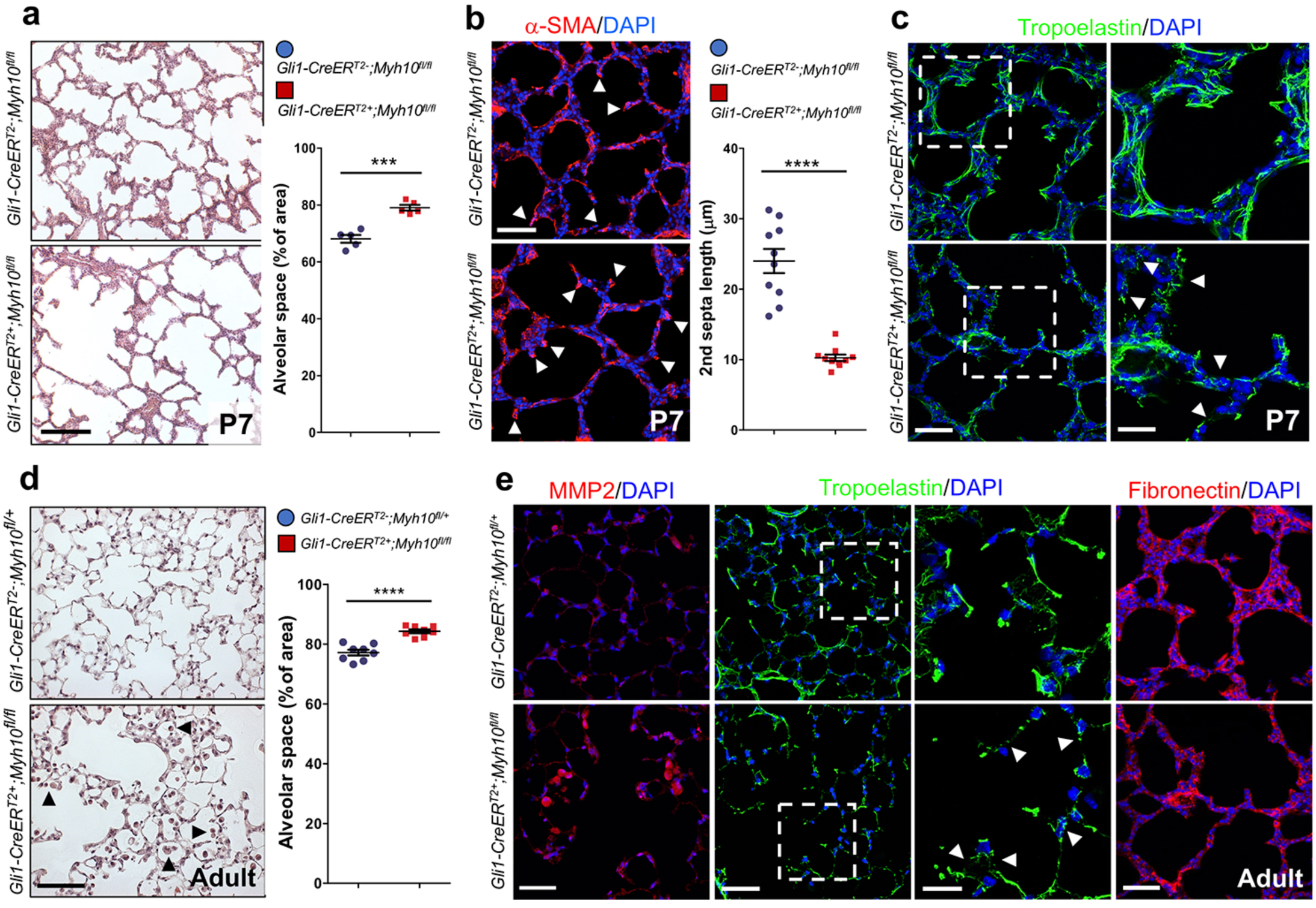
*Myh10* deficiency causes emphysema-like phenotypes. **(a)** H&E staining of wild-type (n=10) and *Gli1-Myh10*^*cKO*^ (n=8) P7 lungs. Quantification of alveolar space in wild-type and *Gli1-Myh10*^*cKO*^ P7 lungs. **(b)** Immunostaining for α-SMA in wild-type (n=10) and *Gli1-Myh10*^*cKO*^ (n=8) P7 lungs. Arrows point to secondary septae. Quantification of secondary septae length in wild-type and *Gli1-Myh10*^*cKO*^ P7 lungs. **(c)** Immunostaining for Tropoelastin in wild-type (n=10) and *Gli1-Myh10*^*cKO*^ (n=8) P7 lungs. Arrowheads point to truncated elastin fibers. **(d)** H&E staining and morphometric analysis of alveolar space in wild-type (n=8) and *Gli1-Myh10*^*cKO*^ (n=6) adult lungs. Arrowheads point to macrophages. **(e)** Immunostaining for MMP2, Tropoelastin and Fibronectin in wild-type (n=8) and *Gli1-Myh10*^*cKO*^ (n=6) adult lungs. Arrowheads point to truncated elastin fibers. Error bars are means ±s.e.m. \*\*\**P* < 0.001, \*\*\*\**P* < 0.0001, two-tailed Student’s *t* test. Scale bars: 100 μm (f (top)), 50 μm (a, d), 30 μm (b, c (left), e (left), f (bottom)), 15 μm (c (right), e (right)).

To test whether *Myh10* participates in lung homeostasis, we deleted *Myh10* in the mesenchymal tissue of adult mice (Supplementary Fig. 7g). *Gli1*-*Myh10*^*cKO*^;*tdTomato* mice were injected with tamoxifen at 2 months, and tdTomato expression was detected in lung mesenchymal cells 1 month later, consistent with previous observations^48^ (Supplementary Fig. 7g, h). At this stage, *Gli1*-*Myh10*^*cKO*^ mice exhibited disrupted septae and enlarged alveolar spaces, accompanied by macrophage accumulation in the alveolar interstitium and alveolar spaces (Fig. 6d and Supplementary Fig. 7i). Moreover, slightly increased numbers of apoptotic cells were observed in *Gli1*-*Myh10*^*cKO*^ lungs compared to wild-type siblings (Supplementary Fig. 7j). The expression of MMP2 was significantly increased in alveolar interstitial tissues as well as in macrophages in *Gli1*-*Myh10*^*cKO*^ lungs compared to wild-type siblings (Fig. 6e), suggesting that upregulated MMP2 in the alveolar interstitium and in macrophages, as well an increased number of macrophages, might be involved in alveolar simplification of *Myh10* deficient adult lungs. Whereas wild-type lungs displayed a well-organized elastin network, *Gli1*-*Myh10*^*cKO*^ lungs showed decreased elastin deposition and truncated and disorganized elastin fibers (Fig. 6e), suggesting that *Myh10* continues to be required to maintain the elastin fiber network and alveolar structure during lung homeostasis.

### MYH10 is down-regulated in emphysematous lungs

Since the conditional knockout phenotype in adult mice is histologically reminiscent of emphysema, we first measured the expression levels of MYH10 protein in human lung samples from healthy donors and emphysema patients. We found significantly downregulated MYH10 expression in lungs of emphysema patients when compared with healthy donor lungs (Fig. 7a, b). Next, we characterized MYH10 expressing cells in human lungs by immunostaining. Similar to MYH10 expression in mouse embryonic and adult lungs, MYH10-expressing human cells did not overlap with NKX2-1-expressing epithelial cells in the alveolar interstitium (Fig. 7c). MYH10 was strongly detected in smooth muscle cells surrounding blood vessels (Fig. 7d). We next analyzed MYH10 expression in human lung samples from 13 patients with emphysema and 7 healthy donors. In healthy lungs, MYH10 was expressed in smooth muscle cells and mesenchymal cells of the alveolar interstitium. In contrast, in emphysematous lungs, MYH10 levels were strongly reduced in smooth muscle cells and moderately reduced in the alveolar interstitium (Fig. 7e, Supplementary Fig. 8a, b). Overall, MYH10 was downregulated in 10 (77%) of the 13 emphysematous specimens (Supplementary Table 1). Taken together, these data indicate that *MYH10* not only has a critical function in lung alveologenesis, but that it may also play a protective role against the occurrence of emphysema.

**Figure 7.**
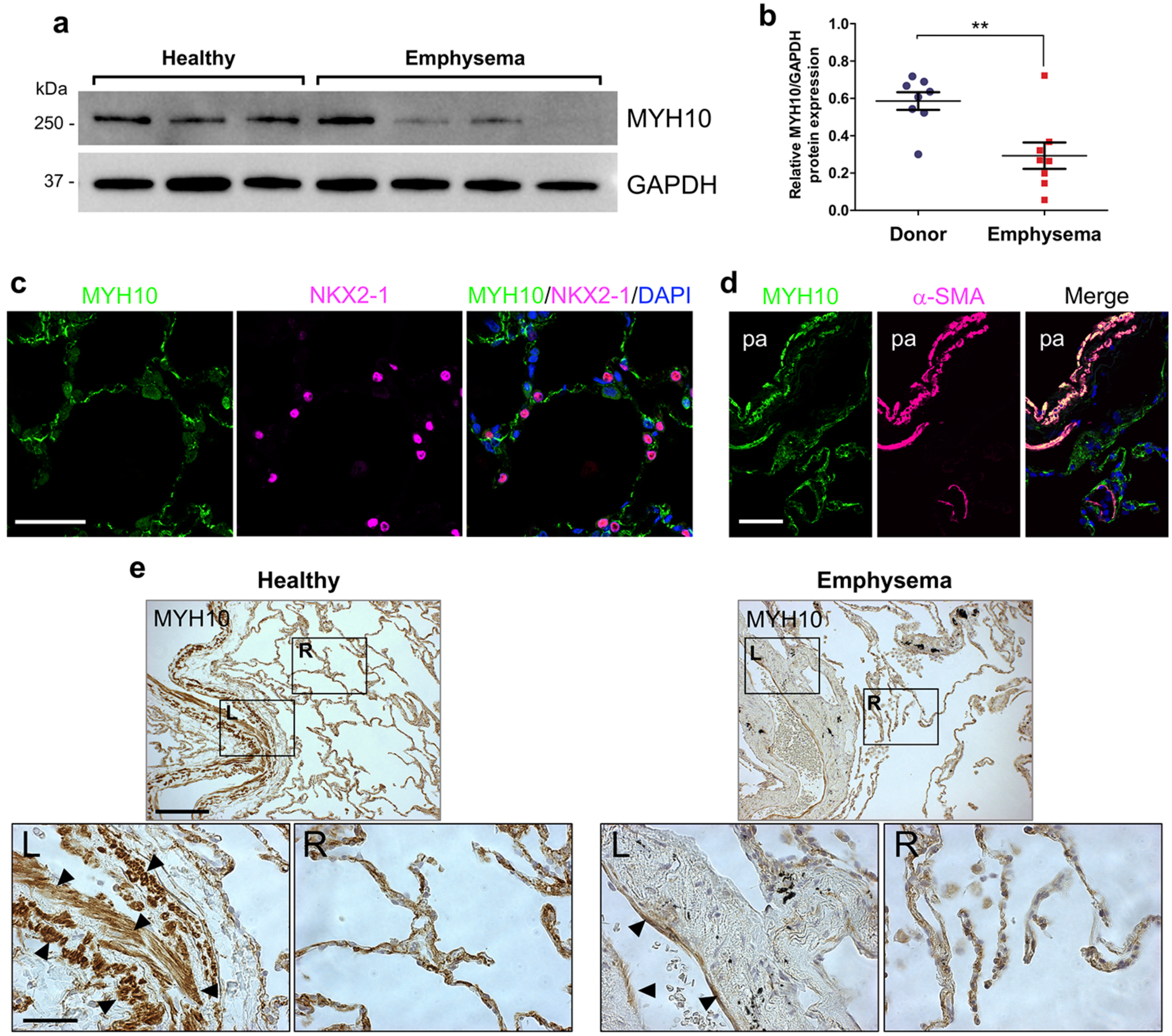
MYH10 is upregulated in emphysematous human lungs. **(a)** Representative western blot for human MYH10 of lung tissues from healthy donors (three individuals) and emphysema patients (four individuals). **(b)** Quantification of MYH10 expression of lung tissues from healthy donors (three individuals) and emphysema patients (four individuals). **(c)** Immunostaining for MYH10 and NKX2-1 in adult human lungs. **(d)** Immunostaining for MYH10 and α-SMA in adult human lungs. pa, pulmonary artery. **(e)** Immunostaining for MYH10 in healthy control and emphysema patient lungs. Arrowheads point to smooth muscle cells. L, left; R, right. Error bars are means ±s.e.m. \*\**P* < 0.01,two-tailed Student’s *t* test. Scale bars: 100 μm (e (top)), 50 μm (c, d), 30 μm (e (bottom)).

## Discussion

Here, we identified a mutation in the mouse *Myh10/NM II-B* gene through an ENU mutagenesis screen and determined its role during lung development and alveologenesis, through a molecular mechanism involving MMP-mediated ECM remodeling. *Myh10* mutant lungs display defects in: (1) ECM remodeling and cell proliferation during the canalicular stage, and (2) distal epithelial cell differentiation, AT1 cell morphogenesis, and interstitial remodeling/tissue morphogenesis during the saccular stage. As a consequence, mutant pups exhibit alveolar collapse and respiratory distress, and die shortly after birth (Supplementary Fig. 9a).

Our results implicate *Myh10* in the maintenance of lung ECM components. The ECM is a three-dimensional, dynamic structure that is constantly remodeled to control tissue homeostasis. Dynamic remodeling of the ECM is essential for organ development and homeostasis and its dysregulation can result in pathological conditions. Thus, proteins involved in ECM degradation and modification, including MMPs, are essential for metazoan development^35,36^. ECM-remodeling MMPs are tightly regulated at the transcriptional, post-transcriptional, and post-translational levels. Examples of biological processes regulating MMP expression and/or function include mechanical forces and actomyosin contractility. Based on our mass spectrometry-based proteomic analysis, we propose a Myosin-dependent regulation of MMP-2 which involves another ECM molecule, THBS. Secretion of THBSs and their distribution in the ECM has been shown to be regulated by the actin cytoskeleton. Elevated ECM deposition of THBS1 or THBS2 is associated with increased tissue stiffness, such as in fibrosis and atherosclerotic plaques^49,50^. Interestingly, in these contexts, intracellular tensile forces (contractility) are also increased^50^, suggesting a reciprocal system by which THBS can modulate the actomyosin network. Our results implicate *Myh10* in the maintenance of lung ECM components. The structural actomyosin network defects caused by the loss of *Myh10*^51,52^ is likely to inhibit the secretion of THBS, in turn leading to de-repression of MMP activity, and disrupted ECM remodeling (Supplementary Fig. 9b). Thus, we propose a signaling axis controlling ECM homeostasis through the actomyosin network, Thrombospondins and Matrix metalloproteinases. Although our data point to a critical role for MYH10/THBS/MMP signaling in ECM remodeling during lung development and homeostasis, how the actomyosin network regulates extracellular deposition of THBS via MYH10 function will require further investigation.

Our study also uncovers a potential link between loss of MYH10 and Chronic Obstructive Pulmonary Disease (COPD). COPD is a common disease worldwide, associated with high morbidity, and characterized by progressive airflow obstruction that is poorly reversible, airway inflammation, and systemic effects^15,16^. The main risk factor for the development of COPD is smoking, but other factors such as air pollutants and genetic determinants have been identified. Pulmonary emphysema, a key symptom of COPD, results from the enzymatic destruction of lung ECM components including elastin and collagen, thereby decreasing tissue stiffness and stability^53-55^. Tissue stiffness has been shown to induce actin cytoskeleton reorganization and contractility, which in turn induces matrix rigidity, and modulates cell adhesion, migration, division and differentiation^56,57^. In our study, we observed that downregulation of MYH10 expression in human emphysema correlates with disease progression, and *Myh10* deficient mice display emphysematous lungs, supporting the notion that *Myh10* deficiency might lead to progressive destruction of the lung mesenchyme. Moreover, two genome-wide studies have reported mutations and reduced expression of two actomyosin-related genes, *Myosin ID* (*MYO1D*)^58^ and *Myosin Light Chain Kinase* (*MYLK*)^59^ in emphysematous human lungs, thus strengthening the relevance of our model.

Excess MMP activity in lungs is known to be associated with pulmonary diseases including COPD, acute respiratory distress syndrome (ARDS), sarcoidosis, and tuberculosis^60,61^. In COPD, MMPs are upregulated and their inhibition prevents disease progression in animal models^62,63^. In the present study of *Myh10* deficiency during lung development and homeostasis, excess MMP2 activity and consequent defective ECM remodeling partially recapitulate the pathophysiological features of human COPD, especially emphysema. Although mutations in the *MYH10* locus have not yet been implicated in the pathogenesis of COPD, further studies are required to investigate possible correlations between NMII/MYH10 and COPD pathology. Our results from animal models and human patients suggest that structural alterations of the actomyosin network by loss of *Myh10* function contribute to the pathogenesis of emphysema. We propose MYH10 as a diagnostic tool for COPD and introduce a mouse model to enable further investigation into the etiology of, and therapeutic approaches for, this disease.

## Methods

### Mouse ENU mutagenesis

ENU-injected C57BL/6 males were obtained from Dr. Monica Justice (Baylor College of Medicine, Houston, TX) ^64^ and crossed with C57BL/6 females to generate G1 males (N=170). G1 males were subsequently crossed with C57BL/6 females to generate G2 females, which were backcrossed to G1 males to generate G3 pups. G3 litters were screened for lung and tracheal malformations from P0 to P7^65^. Genomic DNA from mutant mice and their littermates was isolated for whole-exome sequencing.

### Mouse strains

The following mouse lines were kindly provided for our study: *Myh10*-flox by Dr. Robert Adelstein; *Shh-Cre* and *Gli1-CreER*^*T2*^;*ROSA26*^*tdTomato*^by Dr. Saverio Bellusci; *Tek-CreER*^*T2*^ by Dr. Stefan Offermanns; and *CMV-Cre* by Dr. Thomas Braun. Tamoxifen (Sigma) was dissolved in corn oil at 50 mg/ml and intraperitoneally injected at the indicated developmental stages and frequencies. GM6001 (TOCRIS, Catalog No.:2983) was dissolved in DMSO, and fresh dilutions in saline to a total volume of 200 μl were prepared immediately before administration to pregnant females. Intraperitoneal injection of GM6001 (100 mg/kg) was performed once daily from E16.5 and lung tissues were collected from E18.5 embryos (Fig. 3e). All animal care and use procedures in this study were approved by the local animal ethics committee (Regierungspräsidium Darmstadt, Hessen, Germany).

### Whole-exome sequencing and genotyping

For whole-exome sequencing, genomic DNA samples were extracted from two wild-type and two mutant pups according to standard protocols. The whole-exome sequencing was performed by the Beijing Genomics Institute (BGI). Genomic DNA was captured using Agilent SureSelect Biotinylated RNA Library (BAITS) and loaded on a HiSeq2000 platform for sequencing. Sequence reads were aligned to the C57BL/6J mouse reference genome (mm10) and analyzed using CLCBio Genomic Workbench and GATK software. SNPs were called using the GATK haplotype caller, closely following the GATK best practices. Briefly, aligned reads were de-duplicated and re-aligned to all exonic sequences from the Gencode genome annotation (version vM4), taking into account known variants from DBsnp. After base re-calibration, SNP calling was performed within exons, allowing a padding of 100 bp into flanking introns. Variant calls were annotated to known SNPs from DBsnp (version 142) and the functional relevance of variant calls was predicted using Annovar. Next, calls from wild-type mice were merged (unified), while calls from mutant mice were intersected. Intersecting calls were classified as mutant/homozygous if the corresponding wild-type call was heterozygous/absent or as mutant/heterozygous if there was no corresponding wild-type call. For genotyping, an Eco Real-Time PCR System (Illumina) was used for PCR reactions and high-resolution melt analysis. PCR reaction conditions were as follows: 95 °C for 15 s, 40 cycles of 95 °C for 2 s, 60 °C for 2 s and 72 °C for 2 s. Following PCR, a high resolution melt curve was generated by collecting SYBR-green fluorescence data in the 65–95 °C range. The analyses were performed on normalized derivative plots. The following primers were used: *Myh10* (Exon 12) forward 5’-TGG ATA GGA CCA AAC GCC AG -3’ and *Myh10* (Exon 12) reverse 5’-GGA GAG AGT TGA GGA ATA CCT C -3’

### Plasmids

Total RNA and cDNA were obtained from E13.5 lungs using TRIzol reagent (Life technologies) and Superscript III reverse transcriptase (Life technologies), respectively, according to manufacturer’s instructions. cDNA encoding the MYH10 (NCBI Acc. No. NM_175260) wild-type and MYH10^L458R^ proteins was PCR-amplified using total cDNA from E13.5 lungs as a template. PCR fragments were cloned into the pcDNA3.1-Myc-His vector at the HindIII site using the Cold Fusion Cloning Kit (System Biosciences). *Thbs2* (NCBI Acc. No. NM_011581) cDNA was PCR-amplified using total cDNA from E18.5 lungs as a template. PCR fragments were ligated to the pcDNA3.1-Myc-His vector at XhoI/EcoRI sites. The following primers were used: *Myh10* forward 5’-TCC GAG CTC GGT ACC ACT GTT TAC AAT GGC CCA GAG-3’ and *Myh10* reverse 5’-TTG TTC GGG CCC AAG CTC TGA TTG GGG TGG CTG TG-3’; *Thbs2* forward 5’-AGG TCG ACG TCA CAG GTG GAG ACA AGA TG -3’ and *Thbs2* reverse 5’-TGG AAT TCT CGA GGC ATC TCT GCA CTC A -3’. To synthesize RNA probes for *in situ* hybridization, cDNA was PCR-amplified using total cDNA from E13.5 lungs as a template. PCR fragments were cloned into the pGEM T-easy vector (Promega). The following primers were used: *Myh10* forward 5’-GTA CAG AAA GCC CAG ACC AAA GA-3’ and *Myh10* reverse 5’-TTC TTC ATC AGC CAC TCA TCT GC-3’; *Tbx4* forward 5’-CTT CTA CCA CTG CCT GAA GCG T-3’ and *Tbx4* reverse 5’-AGT CTC GTC ATC CAT CGG TCC A-3’.

### *In situ* hybridization and immunohistochemistry

*In situ* hybridization was performed according to standard procedures. Digoxigenin-labelled RNA probes for *Myh10* and *Tbx4* (NCBI Acc. No. NM_011536) were used. Hybridization and probe washes were performed at 70 °C. Immunohistochemistry was performed according to a standard protocol. The following antibodies and dilutions were used for immunostaining: Acetylated α-tubulin (Sigma, 1:5000), CGRP (Abcam, 1:5000), Cleaved Caspase 3 (Cell signaling, 1:500), c-Myc (Santa Cruz Biotechnology, 1:500), Col I (Abcam, 1:500), Col II (Santa Cruz Biotechnology, 1:100), Col IV (Abcam, 1:500), E-cadherin (Santa Cruz Biotechnology, 1:500), Fibronectin (Abcam, 1:1000), HOPX (Santa Cruz Biotechnology, 1:200), Isolectin B4, FITC-conjugate/IsoB4 (Sigma, 1:500), Laminin (Sigma, 1:1000), LAMP3 (1:100, Imgenex), MMP2 (Millipore, 1:500), MYC (Santa Cruz Biotechnology (9E10), 1:1000), MYH-10 (Sigma, 1:500), NG2 (Millipore, 1:500), NKX2-1 (Santa Cruz Biotechnology, 1:200), PCNA (DAKO, 1:200), PDGFR-β (R&D Systems, 1:200), PDPN/T1α (Developmental Studies Hybridoma Bank, 1:20), PECAM/CD31 (Developmental Studies Hybridoma Bank, 1:1), Phalloidin, Rhodamine-conjugate (Thermo Fisher, 1:1000), Phospho-Histone H3 (Cell Signaling, 1:1000), RAGE (R&D Systems, 1:200), SCGB1A1/CC10 (Millipore, 1:1000), SFTPC/ProSP-C (Millipore, 1:1000), Smooth muscle actin/α-SMA (Sigma, 1:1000), THBS1 (Santa Cruz Biotechnology, 1:200), THBS2 (Santa Cruz Biotechnology, 1:200), Tropoelastin (Abcam, 1:1000), and VE-cadherin (BD Biosciences, 1:500).

### RT-qPCR and Western blot analysis

Total RNA and cDNA were obtained from fetal and postnatal lungs using TRIzol reagent (Invitrogen) and Superscript III reverse transcriptase (Invitrogen), respectively, according to the manufacturer’s instructions. For quantitative reverse transcription PCR, the CFX Connect Real-Time system (Bio-Rad) and DyNAmo colorFlash SYBR green qPCR kit (ThermoFisher Scientific) were used. The primer sequences used for qPCR are shown in supplemental information Table S2. For western blots, protein extracts were prepared from fetal and postnatal lungs by using NP-40 buffer (150 mM NaCl, 1.0 % NP-40, 50 mM Tris pH 8.0, 1 mM phenylmethane sulfonyl fluoride (PMSF)). Western blotting was performed according to standard protocols. Primary antibodies were incubated overnight at 4 °C in 3% skim milk. The following antibodies were used: α-Actin (Sigma, 1:2000), Col I (1:2000), GAPDH (Cell Signaling, 1:2000), Fibronectin (1:2000), MMP2 (Millipore, 1:2000), MMP9 (Millipore, 1:2000), MT1-MMP/MMP14 (Santa Cruz Biotechnology, 1:500), MYH10 (1:2000), THBS2 (1:500), and Tropoelastin (1:2000).

### Histology and head skeletal staining

Whole lungs were fixed in 4% formaldehyde, embedded in paraffin and sectioned at a thickness of 5 μm. For cryosections, fixed tissues were embedded in OCT and 10 μm thick coronal sections were collected. Hematoxylin and eosin (H&E) and Periodic Acid Schiff (PAS) staining were performed according to standard protocols. For head skeletal staining, newborn mice were deskinned, dehydrated in 95% ethanol for 1 day and acetone for 1 day. Skeletons were stained with 0.03% alcian blue (Sigma) and 0.005% alizarin red (Sigma).After staining, samples were cleared in 2 % potassium hydroxide and transferred to 50% glycerol for 2 days.

### Transmission Electron Microscopy (TEM)

For TEM analysis, tissues were fixed in 3% glutaraldehyde, followed by 4% osmium tetraoxide (OsO4) in 0.1 mM sodium cacodylate buffer (pH 7.4). After dehydration in ethanol and propylene oxide, the samples were embedded in Epon 812. Ultrathin sections were cut with a diamond knife and stained with uranyl acetate and lead citrate, and examined with a JEM1400 electron microscope (JEOL).

### Cell culture, chemical treatments, and transfections

NIH3T3 cells and *Myh10* null mouse embryonic fibroblasts were provided by Dr. Stefan Offermanns and Dr. Robert S. Adelstein, respectively. Embryonic fibroblasts, and NIH3T3 cells were maintained at 37 °C in a 5% CO2 incubator in DMEM, supplemented with 10% fetal bovine serum, 50 g/ml streptomycin, 50 U/ml penicillin, and 250 ng/ml amphotericin B. Cells were treated with DMSO, Blebbistatin (10 μM), GM6001 (10 μM), and MMP2 inhibitor I (Santa Cruz Biotechnology, SC-204092, 10 μM) at 37 °C in a 5% CO2 incubator for 24-48 h. Transfection was performed using Lipofectamine 2000 transfection reagent (ThermoFisher Scientific) according to manufacturer’s instructions.

### MMP activity assays

For gelatin zymography, lung tissues were homogenized in homogenization buffer (50 mM Tris-HCl, 0.5% Triton X-100, pH 7.4) and supernatant containing gelatinases was collected following centrifugation. For the MMP activity assay, a standard commercial assay kit (Abcam) was used to detect the general activity of MMPs in *Myh10* null and wild-type mouse embryonic fibroblasts according to manufacturer’s instructions. Cells were cultured at 37 °C in a 5% CO2 incubator in serum-free DMEM for 48 hr. Cell lysates and conditioned media were incubated with a fluorescent substrate and fluorescence was measured using a fluorescence microplate reader (FLUOstar Omega, BMG LABTECH).

### siRNA Knockdowns

siRNAs for *EGFP* (EHUEGFP, Sigma) and *Mmp2* (EMU04861, Sigma) were transfected using RNAiMAX transfection reagent (13778, Life Technologies) according to manufacturer’s instructions.

### Fibronectin Assembly Assay

*Myh10* null and wild-type mouse embryonic fibroblasts were cultured for 24h in complete growth medium. Cells were then kept for 24h in starvation medium and scraped into 2% Deoxycholate (DOC) lysis buffer. DOC-soluble material was removed, and SDS-PAGE sample buffer was added to these lysates. DOC-insoluble material was resuspended in solubilization buffer and treated with SDS-PAGE sample buffer. All samples were boiled for 5 min before western blot analysis.

### Transcriptomic analysis (RNAseq)

For RNAseq, total RNA was isolated from E17 lungs (left lobes) of two *Myh10*^*+/+*^ and two *Myh10*^*-/-*^ mice using Trizol (Life Technologies) combined with on-column DNase digestion (TURBO DNAse, Ambion) to avoid contamination by genomic DNA. RNA and library preparation integrity were verified with a BioAnalyzer 2100 (Agilent) or LabChip Gx Touch 24 (Perkin Elmer). 3 µg of total RNA was used as input for Truseq Stranded mRNA Library preparation following the low sample protocol (Illumina). Sequencing was performed on a NextSeq500 instrument (Illumina) using v2 chemistry, resulting in 30M – 50M reads per library, with 1x75bp single-end setup. The resulting raw reads were assessed for quality, adapter content and duplication rates with FastQC (http://www.bioinformatics.babraham.ac.uk/projects/fastqc). Reaper version 15-065 was employed to trim reads after a quality drop below a mean of Q20 in a window of 20 nucleotides. Only reads between 15 and 75 nucleotides were considered for further analyses. Trimmed and filtered reads were aligned to the Ensembl mouse genome version mm10 (GRCm38) using STAR 2.4.0a with the parameter “--outFilterMismatchNoverLmax 0.1” to increase the maximum ratio of mismatches to mapped length to 10%. The number of reads aligning to genes was counted with the featureCounts 1.5.1 tool from the Subread package. Only reads mapping at least partially inside exons were admitted and aggregated per gene. Reads overlapping multiple genes or aligning to multiple regions were excluded. Differentially expressed genes (DEGs) were identified using DESeq2 version 1.14.1. Only genes with a minimum fold change of ±2 (log2 ±1), a maximum Benjamini-Hochberg corrected p-value of 0.05, and a minimum combined mean of 5 reads were considered to be significantly differentially expressed. The Ensembl annotation was enriched with UniProt data based on Ensembl gene identifiers (Activities at the Universal Protein Resource (UniProt)). To examine the biological significance of the DEGs, we carried out a Gene Ontology (GO) enrichment analysis using the online tool Database for Annotation, Visualization and Integrated Discovery (DAVID) Bioinformatics Resource v6.8 (https://david.ncifcrf.gov/home.jsp)^66^ <u>(Supplementary Dataset 1).</u>

### Proteomic analysis (Mass Spectrometry) Triplicate analyses were performed as follows

#### Sample preparation

Cells were lysed by heating to 70°C for 10 minutes in SDS buffer (4% SDS in 0.1 M Tris/HCl, pH 7.6). DNA was sheared by sonication and cell debris was removed by centrifugation at 16.000 g for 10 minutes. A colorimetric 660 nm protein assay (Pierce) was used to determine the concentration of solubilized proteins in the resulting supernatants. Proteins were subsequently precipitated by addition of four volumes of acetone and incubation at -20°C overnight, followed by pelleting at 14.000 g for 10 minutes and washing the pellet with 90% acetone. Samples were dried to remove acetone completely and dissolved in urea buffer (6 M urea, 2 M thiourea, 10 mM HEPES, pH 8.0). Enzymatic digestion of proteins was performed by in-solution digestion. Briefly, protein disulfide bonds were reduced with 4 mM dithiothreitol and alkylated with 20 mM iodoacetamide. Proteins were then cleaved using Lys-C (50:1 protein-to-enzyme ratio) (Wako Chemicals GmbH) at room temperature for 3 hours, followed by overnight trypsination (50:1 protein-to-enzyme ratio) (Serva) at room temperature. Peptide labeling by reductive dimethylation was performed as previously described^67^. Peptide concentration in trypsin-digested samples was measured using the Fluorimetric Peptide Assay (Pierce). Samples containing equal amounts of peptides (75 µ g) were subjected to the dimethyl (in-solution) labeling protocol. In brief, peptide N-termini and lysine residues were methylated for 1 hour at RT by formaldehyde-H2 and cyanoborohydride (light, control mouse fibroblasts) and formaldehyde-^13^C–D_2_ and cyanoborodeuteride (heavy, Myh10-KD), respectively (all reagents: Sigma). The reaction was quenched by acidification and differentially labeled samples were mixed in 1:1 ratio. Mixed samples were fractionated using the high pH reversed-phase peptide fractionation kit (Pierce) according to manufacturer’s instructions.

#### Liquid chromatography/tandem mass spectrometry

For mass spectrometry analysis, fractionated peptides were reconstituted in 10 µl of solvent A (0.1% formic acid). Peptides were separated using an UHPLC system (EASY-nLC 1000, ThermoFisher Scientific) and 20 cm in-house packed C18 silica columns (1.9 µm C18 beads, Dr. Maisch GmbH) coupled in line to a QExactive HF orbitrap mass spectrometer (ThermoFisher Scientific) using an electrospray ionization source. The gradient employed used linearly increasing concentrations of solvent B (90% acetonitrile, 1% formic acid) over solvent A (5% acetonitrile, 1% formic acid) from 5% to 30% over 215 min, from 30% to 60%, from 60% to 95% and from 95% to 5% for 5 min each, followed by re-equilibration with 5% of solvent B. The flow rate was set to a constant 400 nl/min. Full MS spectra were acquired for a mass range of 300 to 1750 m/z with a resolution of 60,000 at 200 m/z. The ion injection target was set to 3 x 10^6^ and the maximum injection time limited to 20 ms. Ions were fragmented by higher energy collision dissociation (HCD) using a normalized collision energy of 27, an isolation window width of 2.2 m/z and an ion injection target of 5 x 10^5^ with a maximum injection time of 20 ms. Precursors with unassigned charge state and a charge state of 1 were excluded from selection for fragmentation. The duration of dynamic exclusion was set to 20 sec. Resulting tandem mass spectra (MS/MS) were acquired with a resolution of 15,000 at 200 m/z using data dependent mode with a loop count of 15 (top15).

#### Data analysis

MS raw data were processed by MaxQuant (1.5.6.5) using the Uniprot mouse database (as of 20/04/2017) containing 89316 entries and the following parameters: a maximum of two missed cleavages, mass tolerance of 4.5 ppm for the main search, trypsin as the digesting enzyme, carbamidomethylation of cysteins as a fixed modification and oxidation of methionine as well as acetylation of the protein N-terminus as variable modifications. For protein quantification based on dimethyl labeling, isotope labels were configured for peptide N-termini and lysine residues with a monoisotopic mass increase of 28.0313 and 36.0757 Da for the light and heavy labels, respectively. Peptides with a minimum of seven amino acids and at least one unique peptide were included in protein identification. MaxQuant was instructed to filter for 1% false discovery rate on both the peptide and protein levels. Only proteins with at least two peptides and one unique peptide were considered as identified and were used for further data analysis. The resulting data contained 2,949 protein groups and was subjected to differential expression analysis using the in-house R package autonomics (https://bitbucket.org/graumannlabtools/autonomics; version 1.0.21), which makes heavy use of functionality provided by the limma package^68^. The analysis considered protein groups that were quantified in at least one of the triplicate samples, with lack of completeness strongly penalized by the underlying Bayesian-moderated t-testing. To examine the biological significance of the differentially expressed proteins, we used autonomics to carry out overrepresentation analysis for Gene Ontology (GO) terms using a Fisher exact test (Supplementary Dataset 2).

### Human emphysematous lung samples

Emphysematous lungs and healthy control lungs were provided by the UGMLC Giessen Biobank, which is a member of the DZL Platform Biobanking. All patients were diagnosed with COPD/emphysema by pulmonary function testing, CT scans and pathological examination from resected lung specimens. The severity of COPD was classified according to the Global Initiative on Obstructive Lung Disease (GOLD) staging system and all subjects belonged to GOLD IV. The severity of emphysema was assessed by pathological inspection of histological samples and graded according to the method as previously described^69^. The study protocol was approved by the Ethics Committee of the Justus-Liebig-University School of Medicine (No. 58/2015), and informed consent was obtained in written form from each patient.

### Morphometric analyses

Pictures of H&E-stained distal lungs were acquired using a Zeiss widefield microscope (AxioImager). Alveolar spaces, thickness of alveolar walls, and length of secondary septae were measured using ImageJ (https://imagej.nih.gov/ij/) according to the American Thoracic Society guidelines^70^. A minimum of 5 representative, non-overlapping fields of view from lungs of at least three mice from each group were evaluated.

### Statistical analysis

Data are presented as mean ±s.e.m of at least three biological replicates. Two-tailed Student’s *t*-tests were used to assess significance. A p-value of < 0.05 was considered significant.

## Data availability

RNA sequencing data from this study have been deposited in GEO under accession code GSE119399 (https://www.ncbi.nlm.nih.gov/geo/query/acc.cgi?acc=GSE119399) (Supplementary Dataset 1). Proteomic analysis data are available from the corresponding authors upon request (Supplementary Dataset 2).

## Acknowledgments

We thank Saverio Bellusci for the *Shh-Cre* and *Gli1-CreER*^*T2*^ mice, Nina Wettschureck for the *Tek-CreER*^*T2*^ mice, and Birgit Spiznagel for the *CMV-Cre* mice. We thank Konstantinos Gkatzis, Arica Beisaw, and Viola Graef for critically reading the manuscript. This work was supported by funds from the Max Planck Society to D.Y.R.S.

## Author contributions

H.-T.K. designed and performed experiments, and wrote the manuscript; W.Y., P.P., F.G., B.G. and C.B. contributed to data analysis and experiments; H.-T.K., Y.-J.J. and S.O. designed and performed *in vitro* studies; S.K. performed TEM analysis; A.M.S., A.M.B., J.G. and H.-T.K. performed mass spectrometry and data analysis; S.G. and H.-T.K. performed RNA sequencing and data analysis; J.P. and M.L. performed data analysis for whole exome sequencing; X.M. and R.S.A. provided *Myh10*^*flox*^ mice and *Myh10* KO embryonic fibroblasts; C.R. and A.G. provided human patient samples; D.Y.R.S. conceived the ENU screen, enabled and supervised the project, analyzed data and edited the manuscript. All authors commented on the manuscript.

## References

1. Burri PH. Fetal and postnatal development of the lung. Annu. Rev. Physiol. 46, 617–628 (1984).

2. Cardoso WV, Lu J. Regulation of early lung morphogenesis: questions, facts and controversies. Development 133, 1611–1624 (2006).

3. Warburton D, et al Lung organogenesis. Curr. Top Dev. Biol. 90, 73–158 (2010).

4. Rock JR, Hogan BL. Epithelial progenitor cells in lung development, maintenance, repair, and disease. Annu. Rev. Cell Dev. Biol. 27, 493–512 (2011).

5. Herriges M, Morrisey EE. Lung development: orchestrating the generation and regeneration of a complex organ. Development 141, 502–513 (2014).

6. Branchfield K, Li R, Lungova V, Verheyden JM, McCulley D, Sun X. A three-dimensional study of alveologenesis in mouse lung. Dev. Biol. 409, 429–441 (2016).

7. McCulley D, Wienhold M, Sun X. The pulmonary mesenchyme directs lung development. Curr. Opin. Genet. Dev. 32, 98–105 (2015).

8. Vicente-Manzanares M, Ma X, Adelstein RS, Horwitz AR. Non-muscle myosin II takes centre stage in cell adhesion and migration. Nat. Rev. Mol. Cell Biol. 10, 778–790 (2009).

9. Heissler SM, Manstein DJ. Nonmuscle myosin-2: mix and match. Cell Mol. Life Sci. 70, 1– 21 (2013).

10. Newell-Litwa KA, Horwitz R, Lamers ML. Non-muscle myosin II in disease: mechanisms and therapeutic opportunities. Dis. Model Mech. 8, 1495–1515 (2015).

11. Berg JS, Powell BC, Cheney RE. A millennial myosin census. Molecular Biology of the Cell 12, 780–794 (2001).

12. Conti MA, Adelstein RS. Nonmuscle myosin II moves in new directions. J. Cell Sci. 121, 11–18 (2008).

13. Ma X, Jana SS, Conti MA, Kawamoto S, Claycomb WC, Adelstein RS. Ablation of nonmuscle myosin II-B and II-C reveals a role for nonmuscle myosin II in cardiac myocyte karyokinesis. Mol. Biol. Cell 21, 3952–3962 (2010).

14. Plosa EJ, Gooding KA, Zent R, Prince LS. Nonmuscle myosin II regulation of lung epithelial morphology. Dev. Dyn. 241, 1770–1781 (2012).

15. Decramer M, Janssens W, Miravitlles M. Chronic obstructive pulmonary disease. Lancet 379, 1341–1351 (2012).

16. Rabe KF, Watz H. Chronic obstructive pulmonary disease. Lancet 389, 1931–1940 (2017).

17. Fischer A, du Bois R. Interstitial lung disease in connective tissue disorders (vol 380, pg 689, 2012). Lancet 380, 1148–1148 (2012).

18. Taraseviciene-Stewart L, Voelkel NF. Molecular pathogenesis of emphysema. J. Clin. Invest. 118, 394–402 (2008).

19. Marcelino MY, Fuoco NL, de Faria CA, Kozma Rde L, Marques LF, Ribeiro-Paes JT. Animal models in chronic obstructive pulmonary disease-an overview. Exp. Lung Res. 40, 259–271 (2014).

20. Perez-Rial S, Giron-Martinez A, Peces-Barba G. Animal models of chronic obstructive pulmonary disease. Arch. Bronconeumol. 51, 121–127 (2015).

21. Justice MJ, Noveroske JK, Weber JS, Zheng B, Bradley A. Mouse ENU mutagenesis. Hum. Mol. Genet. 8, 1955–1963 (1999).

22. Tullio AN, et al Nonmuscle myosin II-B is required for normal development of the mouse heart. Proc. Natl. Acad. Sci. U S A 94, 12407–12412 (1997).

23. Barron L, Gharib SA, Duffield JS. Lung Pericytes and Resident Fibroblasts: Busy Multitaskers. Am. J. Pathol. 186, 2519–2531 (2016).

24. Treutlein B, et al Reconstructing lineage hierarchies of the distal lung epithelium using single-cell RNA-seq. Nature 509, 371–375 (2014).

25. Laresgoiti U, et al Lung epithelial tip progenitors integrate glucocorticoid- and STAT3-mediated signals to control progeny fate. Development 143, 3686–3699 (2016).

26. Ridsdale R, Post M. Surfactant lipid synthesis and lamellar body formation in glycogen-laden type II cells. Am. J. Physiol. Lung Cell Mol. Physiol. 287, L743–751 (2004).

27. Takeda K, Kishi H, Ma X, Yu ZX, Adelstein RS. Ablation and mutation of nonmuscle myosin heavy chain II-B results in a defect in cardiac myocyte cytokinesis. Circ. Res. 93, 330–337 (2003).

28. Ma X, Bao J, Adelstein RS. Loss of cell adhesion causes hydrocephalus in nonmuscle myosin II-B-ablated and mutated mice. Mol. Biol. Cell 18, 2305–2312 (2007).

29. Shutova M, Yang C, Vasiliev JM, Svitkina T. Functions of nonmuscle myosin II in assembly of the cellular contractile system. PLoS One 7, e40814 (2012).

30. Zhang W, et al Spatial-temporal targeting of lung-specific mesenchyme by a Tbx4 enhancer. BMC Biol. 11, 111 (2013).

31. Bonnans C, Chou J, Werb Z. Remodelling the extracellular matrix in development and disease. Nat. Rev. Mol. Cell Biol. 15, 786–801 (2014).

32. Rao Y, Hao R, Wang B, Yao TP. A Mec17-Myosin II Effector Axis Coordinates Microtubule Acetylation and Actin Dynamics to Control Primary Cilium Biogenesis. PLoS One 9, e114087 (2014).

33. Hong H, Kim J, Kim J. Myosin heavy chain 10 (MYH10) is required for centriole migration during the biogenesis of primary cilia. Biochem. Biophys. Res. Commun. 461, 180–185 (2015).

34. Cardoso WV. Molecular regulation of lung development. Annu. Rev. Physiol. 63, 471– 494 (2001).

35. Lu PF, Takai K, Weaver VM, Werb Z. Extracellular Matrix Degradation and Remodeling in Development and Disease. Csh Perspect Biol. 3, (2011).

36. Page-McCaw A, Ewald AJ, Werb Z. Matrix metalloproteinases and the regulation of tissue remodelling. Nat. Rev. Mol. Cell Biol. 8, 221–233 (2007).

37. Rodriguez D, Morrison CJ, Overall CM. Matrix metalloproteinases: What do they not do? New substrates and biological roles identified by murine models and proteomics. Bba-Mol. Cell Res. 1803, 39–54 (2010).

38. Greenlee KJ, Werb Z, Kheradmand F. Matrix metalloproteinases in lung: multiple, multifarious, and multifaceted. Physiol. Rev. 87, 69–98 (2007).

39. Lo CM, Buxton DB, Chua GC, Dembo M, Adelstein RS, Wang YL. Nonmuscle myosin IIb is involved in the guidance of fibroblast migration. Mol. Biol. Cell 15, 982–989 (2004).

40. Kalson NS, et al Nonmuscle myosin II powered transport of newly formed collagen fibrils at the plasma membrane. Proc. Natl. Acad. Sci. U S A 110, E4743–4752 (2013).

41. Wang Z, et al Interleukin-lbeta induces migration of rat arterial smooth muscle cells through a mechanism involving increased matrix metalloproteinase-2 activity. J. Surg. Res. 169, 328–336 (2011).

42. Kielty CM, Sherratt MJ, Shuttleworth CA. Elastic fibres. J. Cell Sci. 115, 2817–2828 (2002).

43. Yanagisawa H, Davis EC. Unraveling the mechanism of elastic fiber assembly: The roles of short fibulins. Int. J. Biochem. Cell Biol. 42, 1084–1093 (2010).

44. Adams JC, Lawler J. The thrombospondins. Cold Spring Harb Perspect Biol. 3, a009712 (2011).

45. Calabro NE, Kristofik NJ, Kyriakides TR. Thrombospondin-2 and extracellular matrix assembly. Bba-Gen. Subjects 1840, 2396–2402 (2014).

46. Resovi A, Pinessi D, Chiorino G, Taraboletti G. Current understanding of the thrombospondin-1 interactome. Matrix Biol. 37, 83–91 (2014).

47. Ma X, et al Conditional ablation of nonmuscle myosin II-B delineates heart defects in adult mice. Circ. Res. 105, 1102–1109 (2009).

48. Kramann R, et al Perivascular Gli1+ progenitors are key contributors to injury-induced organ fibrosis. Cell Stem Cell 16, 51–66 (2015).

49. Hellewell AL, Gong XY, Scharich K, Christofidou ED, Adams JC. Modulation of the extracellular matrix patterning of thrombospondins by actin dynamics and thrombospondin oligomer state. Bioscience Rep. 35, (2015).

50. Riessen R, Kearney M, Lawler J, Isner JM. Immunolocalisation of thrombospondin-1 in human atherosclerotic and restenotic arteries. Am. Heart J. 135, 357–364 (1998).

51. Zaidel-Bar R, Guo ZH, Luxenburg C. The contractome - a systems view of actomyosin contractility in non-muscle cells. J. Cell Sci. 128, 2209–2217 (2015).

52. Pandya P, Orgaz JL, Sanz-Moreno V. Actomyosin contractility and collective migration: may the force be with you. Curr. Opin. Cell Biol. 48, 87–96 (2017).

53. Finlay GA, O’Donnell MD, O’Connor CM, Hayes JP, FitzGerald MX. Elastin and collagen remodeling in emphysema. A scanning electron microscopy study. Am. J. Pathol. 149, 1405–1415 (1996).

54. Birukov KG. Balancing between stiff and soft: a life-saving compromise for lung epithelium in lung injury. J. Appl. Physiol. (1985) 117, 1213–1214 (2014).

55. Takahashi A, Majumdar A, Parameswaran H, Bartolak-Suki E, Suki B. Proteoglycans maintain lung stability in an elastase-treated mouse model of emphysema. Am. J. Respir. Cell Mol. Biol. 51, 26–33 (2014).

56. Kumper S, Marshall CJ. ROCK-Driven Actomyosin Contractility Induces Tissue Stiffness and Tumor Growth. Cancer Cell 19, 695–697 (2011).

57. Humphrey JD, Dufresne ER, Schwartz MA. Mechanotransduction and extracellular matrix homeostasis. Nat. Rev. Mol. Cell Biol. 15, 802–812 (2014).

58. Castaldi PJ, et al Genome-wide association identifies regulatory Loci associated with distinct local histogram emphysema patterns. Am. J. Respir. Crit. Care Med. 190, 399–409 (2014).

59. Golpon HA, et al Emphysema lung tissue gene expression profiling. Am. J. Respir. Cell Mol. Biol. 31, 595–600 (2004).

60. Elkington PT, Friedland JS. Matrix metalloproteinases in destructive pulmonary pathology. Thorax 61, 259–266 (2006).

61. Baraldo S, et al Matrix metalloproteinase-2 protein in lung periphery is related to COPD progression. Chest 132, 1733–1740 (2007).

62. Wright JL, Cosio M, Churg A. Animal models of chronic obstructive pulmonary disease. Am. J. Physiol. Lung Cell Mol. Physiol. 295, L1–15 (2008).

63. Navratilova Z, Kolek V, Petrek M. Matrix Metalloproteinases and Their Inhibitors in Chronic Obstructive Pulmonary Disease. Arch Immunol. Ther. Exp. (Warsz) 64, 177–193 (2016).

64. Salinger AP, Justice MJ. Mouse Mutagenesis Using N-Ethyl-N-Nitrosourea (ENU). CSH Protoc. 2008, pdb prot4985 (2008).

65. Yin W, Kim HT, Wang S, Gunawan F, Wang L, Kishimoto K, Zhong H, Roman D, Preussner J, Guenther S, Graef V, Buettner C, Grohmann B, Looso M, Morimoto M, Mardon G, Offermanns S. Stainier DYR. The potassium channel KCNJ13 is essential for smooth muscle cytoskeletal organization during mouse tracheal tubulogenesis. Nat. Commun. 9(1), 2815 (2018).

66. Huang DW, Sherman BT, Lempicki RA. Systematic and integrative analysis of large gene lists using DAVID bioinformatics resources. Nat. Protoc. 4, 44–57 (2009).

67. Boersema PJ, Raijmakers R, Lemeer S, Mohammed S, Heck AJ. Multiplex peptide stable isotope dimethyl labeling for quantitative proteomics. Nat. Protoc. 4, 484–494 (2009).

68. Ritchie ME, Phipson B, Wu D, Hu Y, Law CW, Shi W, Smyth GK. limma powers differential expression analyses for RNA-sequencing and microarray studies. Nucleic Acids Res. 43(7), e47 (2015).

69. Thurlbeck WM, Dunnill MS, Hartung W, Heard BE, Heppleston AG, Ryder RC. A comparison of three methods of measuring emphysema. Hum. Pathol. 1, 215–226 (1970).

70. Hsia CC, Hyde DM, Ochs M, Weibel ER; ATS/ERS Joint Task Force on Quantitative Assessment of Lung Structure. An official research policy statement of the American Thoracic Society/European Respiratory Society: standards for quantitative assessment of lung structure. Am. J. Respir. Crit. Care Med. 181, 394–418 (2010).

